# Functional and structural profiling of circulation via genetically encoded modular fluorescent probes

**DOI:** 10.1101/2025.03.18.643859

**Authors:** Marta Vittani, Ashley Bomin Lee, Xiaowen Wang, Yuichi Hiraoka, Daichi Sasaki, Ayumu Konno, Philip Gade Knak, Peter Kusk, Masaki Nagao, Julien Courtin, Muhammad Fadhli Putranto, Yusuke Nasu, Saki Tsuno, Ken Ueda, Yuri Osuga Yoshiike, Takashi Tsuboi, Alma Piganiol, Thomas Bienvenu, Miho Terunuma, Hirokazu Hirai, Maiken Nedergaard, Kohichi Tanaka, Antonios Asiminas, Hajime Hirase

**Author notes:** contributed equally (Marta Vittani, Ashley Bomin Lee, Xiaowen Wang, Yuichi Hiraoka). Correspondence should be addressed to Hajime Hirase.

## Abstract

Sustained labeling of fluids is crucial for their investigation in animal models. Here, we introduce a mouse line (Alb-mSc-ST), where blood and interstitial fluid are labeled with the red fluorescent protein mScarlet and SpyTag. The SpyTag-SpyCatcher technology is exploited to monitor circulating fluid properties by biosensors or detect blood-brain barrier disruption. This approach represents a valuable tool for studying vascular structure, permeability and microenvironment in body organs *in vivo*.

## Main

Extracellular fluid dynamics in the brain have garnered significant interest due to their role in metabolic waste clearance^1^ and the prevention of protein aggregates associated with neurological diseases^2^. Albumin, a secretory protein produced in the liver^3^, is the most abundant protein in both blood plasma and cerebrospinal fluid (CSF) with a concentration of ∼ 450 µM and ∼ 2 µM, respectively^4,5^. We previously demonstrated that systemic injection of liver-targeted adeno-associated viral vectors (AAVs) enables the expression of fluorescent protein-tagged albumin, allowing for long-term imaging of blood microcirculation^6,7^. Here, we present a novel transgenic mouse model in which mScarlet (mSc), a bright red fluorescent protein^8^, and SpyTag (ST), a peptide tag for covalent protein labeling^9^, are knocked into the albumin locus. This modification generates fluorescent albumin (Alb-mSc-ST), providing a genetic tool for tracking albumin dynamics in vivo (**Fig. 1a−b**).

**Figure 1.**
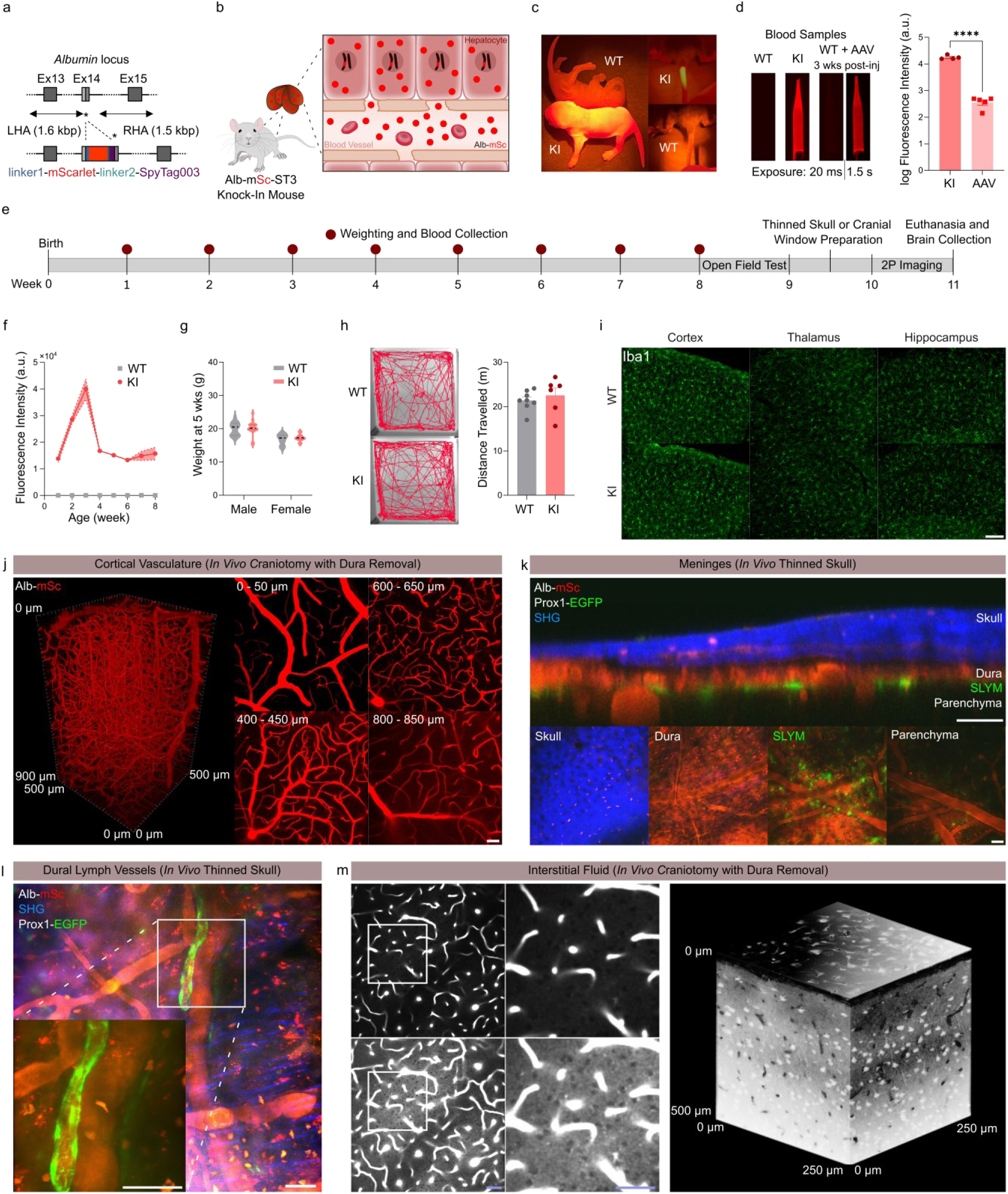
Alb-mSc-ST mice exhibit vascular and interstitial fluid suitable for *in vivo* tissue cytoarchitecture imaging. (a) Genomic map of Alb-mSc-ST knock-in locus. mScarlet and SpyTag003 were knocked into exon 14 of the albumin gene. (b) In heterozygous Alb-mSc-ST mice, Alb-mSc-ST is produced in hepatocytes secreted into liver blood vessels through fenestrations. (c) Optogenotyping of Alb-mSc-ST mice. (d) Blood samples collected from Alb-mSc-ST knock-in (KI) mice, wild-type (WT) littermates and mice injected with AAV8/P3-Alb-mScarlet 3 weeks before collection (left) and their fluorescence intensities (right) with y-axis in log_10_ scale (n = 4−5 mice). Unpaired t-test: ****p < 0.0001. (e) Experiment timeline. (f) Blood plasma fluorescence intensity in the first 8 weeks after birth of KI and WT littermates (n = 6−8 mice). (g) Weights of KI and WT littermates at 5 weeks of age (n = 7−19 mice per group). Mixed-effects analysis using a REML model: no significant effect of genotype (F_1, 52_ = 0.5178, p = 0.475). (h) Example traces of mouse trajectory (left) and total distance traveled (right) for the last 6 min of 10 min recording (n = 6−8 mice). Unpaired t-test: p = 0.5076. (i) Example images of brain slices of KI and WT littermates immunostained for microglia by Iba1. Scale bar, 100 µm. (j) Volumetric imaging of brain vasculature covering 950 µm below the pial surface of Alb-mSc-ST mouse (left) and representative images at various depth (right). Scale bar, 50 µm. (k) Cross-section of thinned skull and meninges from an Alb-mSc-ST x Prox1-EGFP^+^ mouse (top) and representative images of various subcranial structures (bottom). Scale bar, 50 µm. (l) Representative image of dural blood vessels (red), lymph vessels (green) and collagen fibers (blue) in an Alb-mSc-ST x Prox1-EGFP^+^ mouse. Scale bar, 50 µm. (m) Representative image of brain parenchyma of Alb-mSc-ST mouse with normal (left, top) and high (left, bottom) contrast adjustment, the latter displaying dark spots for cellular elements. Volumetric reconstruction with an inverted scale, visualizes the neural cytoarchitecture (right). Scale bar, 50 µm. All graphs show means ± SEM.

Heterozygous Alb-mSc-ST mice were easily identifiable from their skin fluorescence under a green LED flashlight (**Fig. 1c**), reflecting the presence of fluorescent albumin in the dermis. Alb-mSc-ST mice exhibited normal physiology and behavior, with no difference in body weight and ambulatory activity compared to wild-type littermates in open field test (**Fig. 1g−h, S1a−c, e**). Microglial morphology remained unchanged, with no signs of reactivity, suggesting that hemizygous replacement of *Alb* by *Alb-mSc-ST* does not induce brain inflammation (**Fig. 1i**). Fluorescent albumin was robustly present in plasma at birth, with estimated concentrations of ∼ 200 µM (**Fig. 1e−f**). The fluorescence signal was at least an order of magnitude stronger than AAV-expressed Alb-mSc (AAV8/P3-Alb-mSc, 2 x 10^11^ vg)^6^ (**Fig. 1d**), with no detectable sex differences (**Fig. S1d**). The strong plasma fluorescence enables high-resolution vasculature imaging, allowing visualization of blood flow up to ∼1-mm deep - spanning the entire cortex and reaching the white matter - via two-photon microscopy (**Fig. 1j**). High-frame rate imaging captured the capillary-level blood flow dynamics (**Suppl. Video 1**), demonstrating the utility of this model for studying cerebrovascular function in vivo.

Leveraging the interstitial presence of Alb-mSc-ST in peripheral tissues (**Suppl. Video 2**), we sought to visualize the cytoarchitecture of the cerebral meninges. To achieve this, Alb-mSc-ST mice were crossed with lymphatic-specific Prox1-eGFP^+^ mice, and the *in vivo* subcranial structures were imaged using two-photon microscopy through a thinned skull preparation (**Fig. 1k**). Collagen fibers within the skull and dura mater were visualized by second harmonic generation (**Fig. 1k−l**). Additionally, skull osteocytes and dural macrophages exhibited red fluorescence, likely due to Alb-mSc-ST uptake from bone tissue and extracellular space (ECS). The dural structure was revealed by “shadow imaging”, where the bright Alb-mSc-ST background contrasted the dark cellular components (**Fig. 1k**). Albumin fluorescence was also detected in dural lymph (**Fig. 1l, Suppl. Video 3**), while the subarachnoid space flanking the eGFP^+^ cells^10^, showed a reduced signal, consistent with the low protein content of CSF (**Fig. 1k**). CSF was labeled in the perivascular space surrounding large pial vessels (**Fig. S1f**). Shadow imaging extended to the cortical parenchyma through a dura-removed cranial window, clearly distinguishing neurons and dendritic structures (**Fig. 1m**). Deeper brain regions, such as the nucleus accumbens and prefrontal cortex, were visualized using an implantable miniscope (**Suppl. Videos 4 and 5**).

ST forms irreversible covalent bonds with the SpyCatcher (SC) protein^9^. To exploit this interaction, we expressed secretory SC-mNeonGreen (SC-mNG) in liver hepatocytes of Alb-mSc-ST mice, enabling the formation of Alb-mSc-ST and SC-mNG conjugate via ST-SC interaction (**Fig. 2a**). Blood samples from mice systemically injected with AAV8/P3-mNG or AAV8/P3-SC-mNG exhibited a four-orders-of-magnitude increase in plasma green fluorescence signal with SC expression (**Fig. 2b−c, S2a**). Notably, Alb-mSc-ST-SC-mNG fluorescence was threefold brighter than a standard AAV-expressed Alb-mNG (AAV8/P3-Alb-mNG, 2 x 10^11^ vg)^6^ (**Fig. S2c**), demonstrating a superior approach for blood-targeted biosensor expression. Immunoblotting confirmed the formation of the Alb-mSc-ST-SC-mNG macromolecule as expected (**Fig. 2d**), and two-photon imaging validated successful cortical vasculature labeling (**Fig. 2e**). Leveraging this ST-SC technology, we developed a blood-targeted biosensor platform by expressing green SC-tagged fluorescent biosensors in hepatocytes and using Alb-mSc-ST as a circulating molecular carrier. Biosensors expression was confirmed by high fluorescence levels in blood samples (**Fig. 2f, S2d**). Two-photon imaging was used to monitor biosensors responses to physiological stimuli, with simultaneous mSc signal acquisition to account for blood volume variability (**Fig. 2b, S2e, S2p**). Using this approach, we examined blood pH fluctuations (baseline pH = ∼ 7.4) in somatosensory cortex pial vessels during ketamine-xylazine anesthesia using the SC-tagged pH sensitive green fluorescent protein Super Ecliptic Phluorin (SC-SEP, pKa = 7.1)^11^. Whisker stimulation (2 min, 100-ms-long pulses, 5 Hz, 20 psi) induced vessel expansion and a transient pH drop, with signal recovery ∼ 100 s post-stimulus (**Fig. 2j−m, S2g−k**). A similar, but attenuated trend was observed with the less pH-sensitive SC-mNG (pKa = 5.7)^12^ (**Fig. 2i−m, S2j−k**). Additionally, the SC-tagged lactate sensor iLACCO2^13,14^ (SC-iLACCO) detected a reduction in blood lactate during anesthesia, which was counteracted by whisker stimulation (**Fig. S2l−o**). SC-iLACCO also successfully detected increased blood lactate levels following intraperitoneal lactate injection (2 mg/g) in awake mice (**Fig. 2g, S2f**). Finally, using the SC-tagged K^+^ sensor GINKO2^15^ (SC-GINKO), we monitored blood K^+^ dynamics, revealing that topical 1 M KCl application at a distant cortical site reversibly increased blood K^+^ in SC-GINKO mice compared to SC-mNG controls, likely reflecting K^+^ absorption and cortical spreading depression (CSD) (**Fig. 2h, S2q**). Optogenetic induction of CSD via neuronal channelrhodopsin2 (ChR2) also increased blood K^+^, indicating systemic K^+^ uptake following CSD (**Fig. 2i, S2r**).

**Figure 2.**
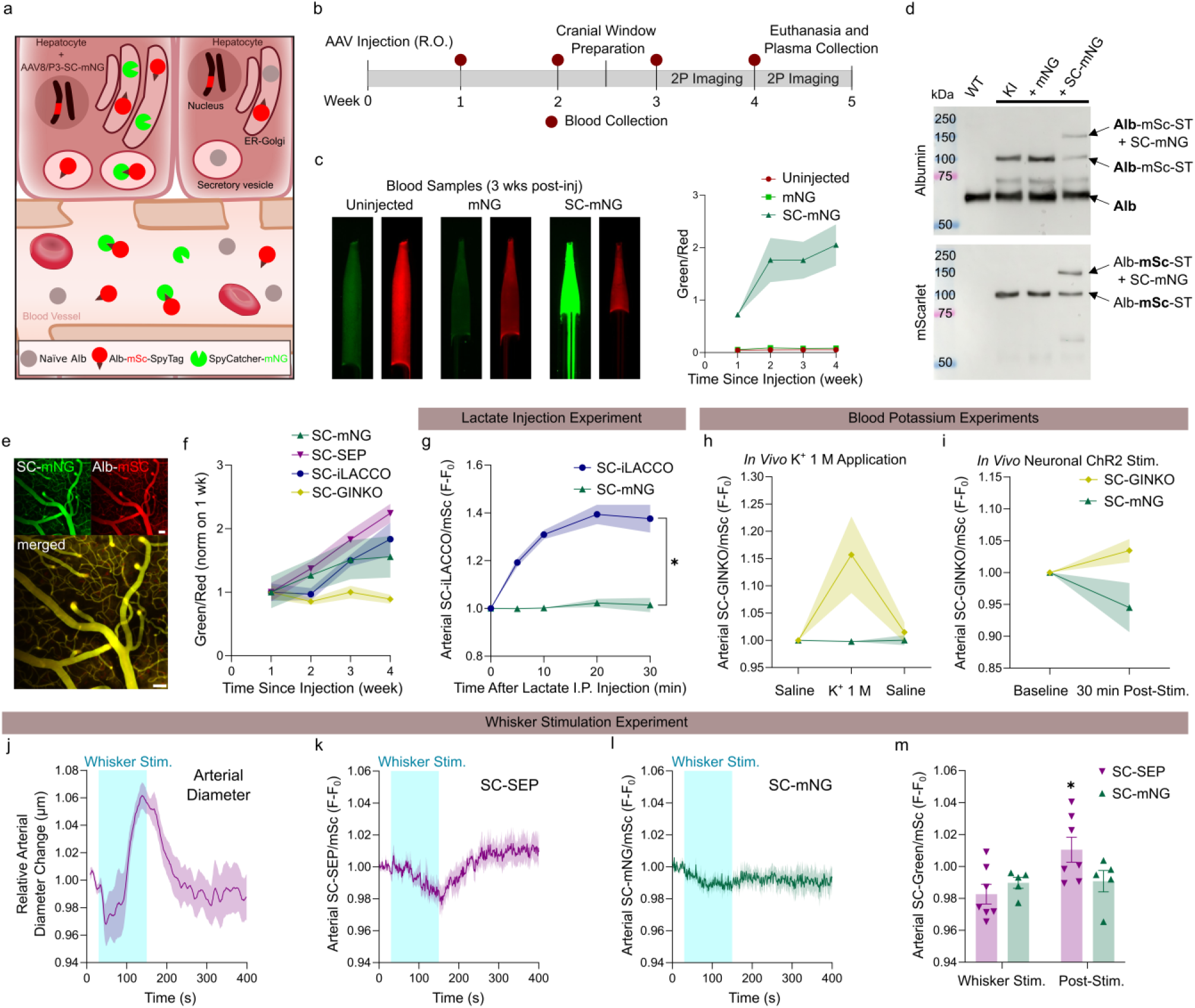
Biosensor attachment to Alb-mSc-ST enables investigation of biochemical properties of blood. (a) Expression of secretory SC-mNG in hepatocytes of Alb-mSc-ST mice leads to irreversible formation of Alb-mSc-ST-SC-mNG through the SC-ST covalent bond. (b) Experiment timeline. (c) Blood samples collected 3 weeks post-injection from Alb-mSc-ST mice injected with AAV8/P3-mNG or AAV8/P3-SC-mNG or uninjected (left). Longitudinal plot of blood green/red fluorescence signal ratio for the first 4 weeks of post-injection (right) (n = 4−5 mice). Two-way ANOVA: significant effect of injection type (F_2, 12_ = 18.22, ***p = 0.0002). (d) Western blot for albumin (top) and mScarlet (bottom) with plasma protein samples from WT littermates and Alb-mSc-ST KI mice injected with AAV8/P3-mNG, AAV8/P3-SC-mNG, or uninjected. (e) Average intensity projection of a 100-μm z-stack showing cortical vasculature of an Alb-mSc-ST mouse injected with AAV8/P3-SC-mNG. Scale bar, 50 µm. (f) Green/red signal ratio in blood samples collected from Alb-mSc-ST mice injected with AAV8/P3-SC-mNG, AAV8/P3-SC-SEP, or AAV8/P3-SC-iLACCO for the first 4 weeks of post-injection (n = 5−7 mice). (g) Green/red signal ratios in cortical arterioles of awake Alb-mSc-ST mice injected with AAV8/P3-SC-iLACCO or AAV8/P3-SC-mNG and imaged after intraperitoneal injection of lactate (2 mg/g, n = 5−6 mice).Two-way ANOVA: significant effect of biosensor (F_1, 9_ = 97.89, ****p < 0.0001). (h) Green/red signal ratios in cortical arterioles of anesthetized Alb-mSc-ST mice injected with AAV8/P3-SC-GINKO or AAV8/P3-SC-mNG following application of either saline or 1 M KCl to a distantcortical site (n = 1−2 recordings from 2−3 mice). Two-way ANOVA: significant effect of applied solution x AAV type interaction (F_2,8_ = 4.726, p = 0.0442). (i) Green/red signal ratios in cortical arterioles of anesthetized Alb-mSc-ST mice injected with AAV8/P3-SC-GINKO or AAV8/P3-SC-mNG following optogenetically induced CSD (n = 1−2 recordings from 2−3 mice). Two-way ANOVA: no significant effect of time x AAV type interaction (F_1, 3_ = 5.785, p = 0.0952). (j−m) Whisker stimulation experiment in anesthetized Alb-mSc-ST mice injected with AAV8/P3-SC-mNG, AAV8/P3-SC-SEP (n = 5−7 mice). Relative diameter change of arterioles in SEP mice (j). Green/red signal ratios in cortical arterioles of SEP and mNG mice (k−l). Quantification of normalized arterial signals (green/red) during and after whisker stimulation (m). Two-way ANOVA: significant effect of stimulation x biosensor interaction (F_1, 10_ = 5.855, *p = 0.0361). All graphs show means ± SEM.

We further leveraged the irreversible ST-SC bond to assess blood brain barrier (BBB) integrity by expressing ECS-facing SC on astrocyte membrane (**Fig. 3a**). To address potential saturation of available SC binding sites, we designed a “destabilized” SC variant by incorporating a PEST sequence^16^ (ECS-dSC) to enhance protein degradation. In vitro validation using HEK293T cells confirmed that ECS-dSC exhibited a higher turnover rate than ECS-SC, making it more suitable for *in vivo* applications (**Fig. 3b**). Since astrocytes enwrap the cerebral vasculature, we selectively expressed ECS-SC or ECS-dSC in astrocytes (**Fig. 3c**). To induce BBB disruption, we applied laser irradiation^17^ or systemic mannitol injection^18^ 40 min before brain collection (**Fig. 3c**). Following laser irradiation, extravasated Alb-mSc-ST was captured by ECS-SC or ECS-dSC, clearly outlining astrocytic endfeet surrounding blood vessels. In contrast, in the absence of ECS-(d)SC expression, Alb-mSc-ST diffused away, confirming the specificity of the interaction (**Fig. 3d**). Histological analysis of laser-irradiated hemisphere revealed mSc fluorescence outlining astrocytic morphology, demonstrating that Alb-mSc-ST successfully binds to ECS-dSC expressing astrocytes and confirming ECS-dSC availability under basal conditions (**Fig. 3e**). Furthermore, the non-irradiated hemisphere of mannitol-injected mice exhibited increased image entropy compared to saline-injected mice, indicating that BBB-permeated Alb-mSc-ST was effectively captured by ECS-dSC and integrated into astrocyte labeling (**Fig. 3f**). Expression of dSC in astrocytes using BBB permeable PhP.eB viral vectors opens the door for brain-wide functional investigation of BBB (**Fig. S3**).

**Figure 3.**
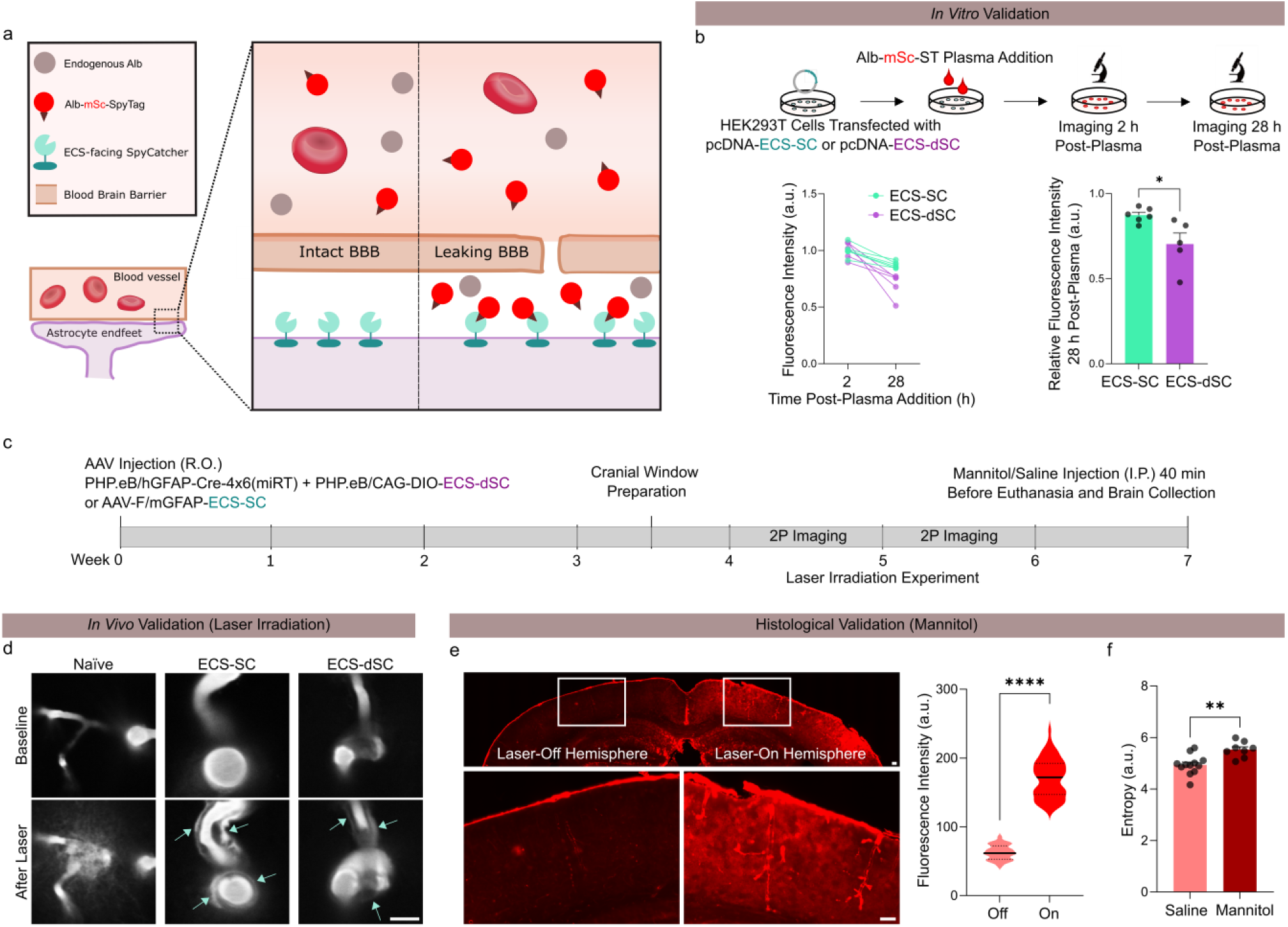
Extravasated Alb-mSc-ST is captured by ECS-facing SC in astrocytes through ST−SC binding. (a) Alb-mSc-ST mice injected with AAVs for astrocyte-targeted expression of ECS-facing SC are used for detecting BBB leakage, as extravasated Alb-mSc-ST will be bound by available ECS-SC at astrocytic endfeet. (b) HEK293T cells expressing either ECS-SC or degradable ECS-dSC were exposed to Alb-mSc-ST plasma and imaged 2h and 28 h post-plasma addition (top). Fluorescence intensity at 2h and 28h (bottom, left) and their ratio (bottom, right) for ECS-SC- and ECS-dSC-expressing cells, showing faster turnover of ECS-dSC. (c) Experiment timeline. (d) Example images of vessels before and after laser irradiation in naïve Alb-mSc-ST mice and Alb-mSc-ST mice expressing ECS-SC or ECS-dSC in astrocytes. Scale bar, 10 µm. (e) Example images of laser-irradiated and contralateral hemispheres in a brain slice from an Alb-mSc-ST mouse expressing astrocytic ECS-dSC (left). Scale bar, 100 µm. Quantification of the fluorescence intensity in the two hemispheres (right). Unpaired t-test: ****p < 0.0001 (n = 3 mice). (f) Entropy levels of cortex and striatum in brain slices from ECS-dSC-expressing Alb-mSc-ST mice injected with the BBB-leakage inducer mannitol or saline. Mann Whitney test: **p = 0.0022 (n = 3 mice, 4 regions each). All graphs show means ± SEM, apart for violin plots in (d, right).

This novel molecular genetic toolset provides a powerful platform for real-time imaging of body fluids and their biochemical properties. In this study, we demonstrated that: (1) perpetual blood plasma labeling with unprecedented brightness enables vascular dynamics imaging from neonate to the aged mice; (2) extracellular albumin fluorescence facilitates high-contrast tissue structural imaging via shadow imaging; (3) plasma functional imaging is achievable through ST-SC system, allowing dynamic biosensor integration for monitoring physiological parameters; (4) BBB integrity can be assessed by expressing ECS-facing SC in astrocytes, enabling precise detection of plasma protein extravasation.

While we primarily explored the utility of Alb-mSc-ST mice the brain, this approach is highly adaptable to a wide range of tissues (**Suppl. Video 2**) and systems biology questions where microcirculation plays a critical role, including cancer^19^ and aging^20^.

## Methods

### Knock-In Mouse

To generate Alb-mScarlet-SpyTag003 mice, we utilized CRISPR/Cas9-mediated genome editing (B6-Alb^em^^1(Alb-mSc-ST3)KTa^, RBRC12137, RIKEN BRC, Japan). The crRNAs targeting exon 14 were designed using the online sgRNA design tool (https://crispr.mit.edu/). We used a crRNA (5’-<u>AUAGUUACCUGAGAAGGUUG</u>guuuuagagcuaugcuguuuug -3’; Fasmac, Kanagawa, Japan, mouse *Alb* gene specific sequence is underlined), which has a double-strand break site in 17 bp downstream of original Alb stop codon and showed the highest digestion efficiency in both *in vivo* and *in vitro* screening assay. We constructed the targeting vector plasmid containing the ScFV linker, mScarlet^8^, and SpyTag003^9^ fused into the carboxyl terminal of mouse Albumin. For homology directed recombination, transgenes were flanked with 1599bp upstream and 1478bp downstream sequences from DSB site, except that the original “TAA” stop codon in the homology arm sequence was changed to “AAA” to make read-through and express following transgenes. The background strain C57BL/6J mice were bought from Japan CLEA, Inc. (Tokyo, Japan). The cryopreserved fertilized pronuclear-stage embryos were prepared by *in vivo* fertilization in human tubal fluid medium (ARK Resource; Kumamoto, Japan) with sperms from two C57BL/6J males and oocytes from ten superovulated females injected with HyperOva (Kyudo, Saga, Japan) and human gonadotropin (ASKA Pharmaceutical, Tokyo, Japan). Next, the targeting plasmid vector (40 ng/µl) and the complex of crRNA (0.61 µM), tracrRNA (0.61 µM), and Cas9 protein (30 ng/µl) (New England Biolab, MA, USA) in nuclease-free PBS (Gibco, MA, USA) were microinjected into the pronucleus of approximately 300 pronuclear-stage embryos. Embryos were then washed and cultured in KSOM medium (ARK Resource) for over an hour. The obtained 245 embryos were transferred to pseudo-pregnant ICR strain mice (Japan CLEA). Fifteen mice were born and prescreened by genomic PCR and sequencing. Three founder generation (F0) mice were verified to have the correct insertion, and two F0 mice were bred with C57BL/6J mice to obtain F1 generation. These two lines backcrossed to C57BL/6J mice at least twice were used for further analyses. We did not discriminate between these two lines in this manuscript because they were indistinguishable in mScarlet fluorescence.

In some experiments (**Fig. 1k−l and S1f**), Alb-mSc-ST knock-in mice were crossed with Prox1-EGFP^+^ reporter mice, which were kindly provided by Dr. Kari Alitalo on C57BL/6JRj background (Janvier Labs, Le Genest-Saint-Isle, France).

Mice were housed in a 12-h light/12-h dark cycle (lights on: 7 am) with food and water *ad libitum*. Optical genotyping was performed by observing ear tissue biopsies using a green LED light (NIGHTSEA Xite-GR). Experimental procedures involving animal care, surgery, and sample preparation performed at the University of Copenhagen were approved by the Danish Animal Experiments Inspectorate and were overseen by the University of Copenhagen Institutional Animal Care and Use Committee (IACUC), in compliance with the European Communities Council Directive of 22 September 2010 (2010/63/EU) legislation governing the protection of animals used for scientific purposes. The procedures for creation and verification of knock-in mice were approved by the Animal Care and Use committee of Tokyo Medical and Dental University.

### AAV Preparation

All codon optimization of the nucleotide sequences and plasmid cloning were performed by Twist bioscience (CA, USA). The nucleotide sequences for the shortened IgK leader (IgKL) signal peptide (M[E]TDTLLLWVLLLWVPGSTGD), the bright green fluorescent protein mNeonGreen (mNG), the SpyCatcher003 (SC) peptide^9^ and the potassium sensor GINKO2^15^ were obtained from the Addgene web site (plasmids #92281,#128144, #133447 and #177116, respectively). The nucleotide sequence for the pH sensor SEP was a codon-optimized version of the sequence from the original publication^11^. Sequence for lactate sensor iLACCO2^13,14^ was a gift from Yusuke Nasu.

For investigating blood microenvironment (**Fig. 2 and S2**), DNA inserts encoding IgKL, SC and a short linker (GSSGS) upstream of the sequence for a biosensor, either SEP or iLACCO or GINKO, or for the control mNG were synthetized and subcloned to the AAV backbone vector pAAV-P3-* ^21^ to construct pAAV-P3-SC-SEP, pAAV-P3-SC-iLACCO, pAAV-P3-SC-GINKO and pAAV-P3-SC-mNG, respectively (Addgene #234541, #234542, #234544 and #234543, respectively). Similarly, a DNA insert encoding IgKL, a short linker upstream of the sequence for mNG was synthesized and subcloned to the same AAV backbone vector to construct pAAV-P3-mNG.

For constructing a transmembrane version of SC facing the extracellular space (ECS), a DNA insert encoding IgKL, SC, a short linker (GSSGS) and a Myc tag (EQKLISEEDL) upstream of the sequence for the PDGFR transmembrane domain was synthesized to obtain the insert named ECS-SC. A second version with higher turnover rate was created by adding the protein degradation tag PEST sequence^16^ after the PDGFR sequence and named ECS-dSC (d = destabilized). For *in vitro* validation of extracellular space-facing SC and dSC (**Fig. 3b, S3a**), the ECS-SC and ECS-dSC inserts were subcloned to the mammalian expression vector pcDNA-* to construct pcDNA-ECS-SC and pcDNA-ECS-dSC. For *in vivo* expression of ECS-SC in astrocytes (**Fig. 3d**), the ECS-SC insert was subcloned to the AAV backbone vector pAAV-mGFAP_*-WPRE-SV40p(A) to construct pAAV-mGFAP-ECS-SC (Addgene #234545). For *in vivo* expression of ECS-dSC in astrocytes (**Fig. 3d−f**), the ECS-dSC insert was subcloned to the AAV backbone vector pAAV-CAG-DIO_*-WPRE-SV40p(A) to construct pAAV-shortCAG-DIO-ECS-dSC (Addgene #234963).

AAV8/P3-mNG (titer: 1.92 x 10^13^ vg/mL), AAV8/P3-Alb-mNG (titer: 3.03 x 10^13^ vg/mL) and AAV-F/mGFAP-ECS-SC (titer: 3.12 x 10^12^ vg/mL) were produced at the Viral Vector Core of Gunma University using the ultracentrifugation method as described previously^22^.

AAV8/P3-SC-SEP (titer: 8.8 x 10^12^ vg/mL), AAV8/P3-SC-iLACCO (titer: 8.9 x 10^12^ vg/mL), AAV8/P3-SC-mNG (titer: 1.0 x 10^13^ vg/mL), AAV8/P3-Alb-mScarlet (titer: 1.7 x 10^13^ vg/mL), PHP.eB/hSyn1-hChR2-mCherry (v124-PHP.eB, titer: 1.4 x 10^13^ vg/mL), AAV-shortCAG-DIO-ECS-dSC (titer: 1.1 x 10^13^ vg/mL) and PHP.eB/hGFAP-Cre-4×6(miRT) (v1013-PHP.eB, titer: 1.4 x 10^13^ vg/mL) were purified by the Viral Vector Facility, Institute of Pharmacology and Toxicology, University of Zurich.

### Cell Culture

HEK293T cells (Dharmacon, HCL4517), cultured in DMEM supplemented with 10% FBS and 50 U/mL penicillin-streptomycin (Thermo Fisher Scientific, 41965039, 16141079 and 15140122), were transfected in a 12-well plate using Fugene HD (Promega, E2311) at 30% confluency. For each well, transfection reagent was mixed with 1 µg of pcDNA-ECS-SC at a 3:1 ratio (mL/mg) in 25 µL Opti-MEM (Thermo Fisher Scientific, 31985070) and added dropwise after 15-minute incubation at room temperature. Each transfection was carried out in two replicates. Both transfected and non-transfected cells (control) were incubated in plasma extracted from Alb-mSc-ST knock-in mice (5 µL per 1 mL culture media) for 10 min and then washed several times with PBS with calcium and magnesium. Cells were imaged using an inverted microscope (Nikon Eclipse Ti) at 2 h and 28 h after transfection.

### AAV Injection

For retro-orbital injection of AAV, the AAV mixture was diluted to 150 µL in sterile saline and administered according to a published protocol^23^. At least 2−3 weeks were allowed for optimal AAV expression before proceeding to further experiments.

For comparison of Alb-mSc expression in knock-in mice versus AAV-injected mice (**Fig. 1d**), wild-type littermates were injected with AAV8/P3-Alb-mScarlet (dosage: 2 x 10^11^ vg) via retro-orbital sinus. For investigating blood microenvironment (**Fig. 2 and S2**), knock-in mice received a retro-orbital injection of either AAV8/P3-SC-SEP (6 x 10^11^ vg) or AAV8/P3-SC-iLACCO (6 x 10^11^ vg) or AAV8/P3-SC-GINKO (6 x 10^11^ vg) or AAV8/P3-SC-mNG (6 x 10^11^ vg) or AAV8/P3-mNG (6 x 10^11^ vg). For comparison of mNG expression (**Fig. S2c**), knock-in mice were injected with AAV8/P3-SC-mNG (6 x 10^11^ vg) and wild-type littermates with AAV8/P3-Alb-mNG (6 x 10^11^ vg) via retro-orbital sinus. For the cortical spreading depression experiment (**Fig. 2i, S2r**), AAV8/P3-SC-GINKO (6 x 10^11^ vg) was coinjected with PHP.eB/hSyn1-hChR2-mCherry (4 x 10^11^ vg) via retro-orbital sinus.

For *in vivo* expression of extracellular space-facing SC or dSC in astrocytes (**Fig. 3d−f**), knock-in mice received a retro-orbital injection of either AAV-F/mGFAP-ECS-SC (6 x 10^11^ vg) or AAV-shortCAG-DIO-ECS-dSC (6 x 10^11^ vg) and PHP.eB/hGFAP-Cre-4×6(miRT) (6 x 10^11^ vg). We estimate that ∼70–80% of cortical areas are represented by ECS-dSC^+^ positive astrocytes.

### *Ex vivo* macro fluorescence imaging

To follow the expression of fluorescent proteins (**Fig. 1, S1, 2 and S2**) over several weeks, blood drops from punctured tail tips of awake mice were collected in borosilicate glass capillaries (1B100F-4 or 1B150F-4, WPI) and examined by a macroscope (Leica M205 FA) equipped with an X-Cite 200Dc light source and a digital camera (C11440 Orca-flash 4.0, Hamamatsu). Filter sets ET GFP (Ex 470/40, Em 525/50, 10447407, Leica) and ET mCherry (Ex 560/40, Em 630/75, 10447407, Leica) were used to image green and red, respectively. Images were acquired using Leica Application Suite X software (version 2.0.0.14332.2).

### Histology

Deeply anesthetized mice (100 mg/kg ketamine, 20 mg/kg xylazine) were transcardially perfused with sterile saline and then with 4% paraformaldehyde (PFA) in 0.1 M phosphate buffer (PB, pH 7.4) using a peristaltic pump. The brain was collected and post-fixed in 4% PFA for 24 h before further storage in PBS. Brain sections of 60 µm were prepared using a vibratome (Leica VT1200 S).

For evaluation of microglia (**Fig. 1i**), brain sections were obtained from knock-in mice. After blocking (5% Normal Goat Serum + PB + 0.1% TritonX-100 for 2 h), the sections were incubated with the primary antibody anti Iba-1 antibody, rabbit (1:1000 for 48 h, 4 °C, 019-19741, Wako). Primary antibody was detected using the secondary antibody anti rabbit-IgG AF488 (1:500 for 2h at RT, A11034, Thermo Fisher Scientific). Finally, stained sections were mounted using ProLongTM Gold antifade regent (P36930, Invitrogen). Images were acquired using Keyence BZ-X710 Fluorescence Microscope and the following settings: 20 X objective lens (Nikon 20x Fluor NA 0.75), exposure 50 ms and GFP filter.

For *ex vivo* validation of ECS-dSC binding extravasated Alb-mSc-ST (**Fig. 3e−f**), brain sections were obtained from knock-in mice injected with AAV-shortCAG-DIO-ECS-dSC and PHP.eB/hGFAP-Cre-4×6(miRT). Sections were mounted using VectaShield Antifade Mounting Medium (H100, Vector Labs). Images were acquired using Keyence BZ-X710 Fluorescence Microscope and the following settings: 20 X objective lens (Nikon 20x Fluor NA 0.75), exposure 50 ms and TxRed filter.

### Plasma Collection

Deeply anesthetized mice (100 mg/kg ketamine, 20 mg/kg xylazine) were intraperitoneally injected with heparin (500U, LEO) 30 minutes before collection to prevent blood coagulation. After exposing the heart and excising the right atrium, the blood was quickly collected with a syringe and transferred in 0.75 mL tubes containing 5 μL EDTA and 5 μL Halt protease and phosphatase inhibitor cocktail (100x, 78429, Thermo Scientific). Blood samples were centrifuged for 10 min at 2000 x g at 4°C. The supernatant, constituting the plasma, was collected and stored in aliquots at −80°C. Plasma aliquots were used for *in vitro* validation of the binding of Alb-mSC-ST to ECS-dSC (**Fig. 3b**).

### Western Blot

The western blot protocol was performed as previously described^24^. Protein samples were subjected to SDS-PAGE gels and transferred to a supported nitrocellulose membrane. Membranes were blocked with blocking buffer (5% bovine serum albumin (BSA) in TBST). Membranes were probed with primary antibodies against albumin (Abcam, ab19194) and RFP (Proteintech, 6G6). Membranes were then probed with HRP-conjugated secondary antibodies and visualized by ECL. Blots were then quantified using the CCD-based FujiFilm LAS 3000 system.

### Open Field Test

The open field arena was a 40 x 40 cm^2^ white foam polyvinyl chloride box, with an inner area of 24 x 24 cm^2^ considered as center zone. Mice were handled daily for 10 min for the 3 days before the experiment. Mice were placed in the behavior room few hours prior testing and tested at the end of the light cycle (5–7 pm). After placing the animal at the corner of the box, the mouse was allowed to explore the arena for 10 min and its movement was recorded with a video camera placed above the arena. Only the last 6 min of the recording were used for analysis. The box was thoroughly cleaned with alcohol and water after each mouse.

### Thinned Skull and Cranial Window Preparation

Throughout the surgery, mice were kept under anesthesia using isoflurane 1.5%. and on a heating pad at 37°C to maintain body temperature. Buprenorphine (0.05 mg/kg) was subcutaneously injected for initial analgesia and lidocaine (0.2 mg/kg) was injected at the site prior to incision for local analgesia. The skull was exposed and a headplate was attached using dental cement (Super Bond C&B, Sun Medical, Shiga, Japan). The area of interest was the somatosensory cortex of the right hemisphere. For thinned skull preparation (**Fig. 1k−l and S1f**), the skull above this area was thinned using a drill until reaching a thickness of ∼ 40−50 µm and a layer of super glue was applied for protection and for inhibiting skull regrowth. For cranial window preparation, a 4-mm diameter craniotomy was drilled, covered with an autoclaved 4-mm or 3-mm diameter coverslip and sealed with dental cement (**Fig. 2 and S2**). For in-depth imaging experiments (**Fig. 1j, 1m**), an intermediate step involving surgical removal of dura mater was added before applying a coverslip. For some SC-GINKO experiments involving induction of CSD (**Fig. 2i, S2r**), two cranial windows were prepared: a 3-mm diameter craniotomy for imaging and a 2-mm diameter craniotomy in the contralateral hemisphere for optogenetic stimulation. For all these preparations, a metal lid was attached with dental silicon for protection. After recovery from anesthesia, the animal was returned to its cage and was injected subcutaneously with the analgesic carprofen (5 mg/kg) immediately after surgery, 24h and 48h post-surgery. For some SC-GINKO experiments (**Fig. 2h, S2q**), on the day of imaging a second acute 2-mm diameter craniotomy was created in the ipsilateral hemisphere and kept open without coverslip for local application of solutions; these mice were euthanized immediately after imaging.

### *In Vivo* Two-Photon Imaging

Two-photon imaging was performed on anesthetized (70 mg/kg ketamine, 10 mg/kg xylazine) mice or on awake mice previously habituated to head-fixation using the following protocol: 30 min on day 1, 1 h on day 2, 1 h on day 3, 1h 30 min on day 4. The Bergamo microscope (Thorlabs) is equipped with a resonant scanner, a Chameleon Ultra 2 laser (Coherent), an objective lens (XLPlan N; ×25 NA = 1.05; Olympus) and the primary dichroic mirror FF705-Di01-25×36 (Chroma). Emission light was separated by the secondary dichroic mirror (FF562-Di03, Semrock) with band-pass filters FF03-525/50 (Semrock) for mScarlet, FF01-607/70 (Semrock) for mNG, SEP and iLACCO, BP447/60 (Semrock) for second harmonic generation (SHG). Excitation wavelength varied according to the experiment: 910 nm for second harmonic generation to visualize collagen fibers (**Fig. 1k, 1l**); 930 nm for simultaneous imaging of mScarlet and EGFP (**Fig. 1k, 1l**); 940 nm for simultaneous imaging of mScarlet and green biosensors (**Fig. 2 and S2**); 1040−1060 nm for mScarlet (**Fig. 1j, 1m**). Of note, mScarlet is visible at both optimal (1060 nm, 90% excitation in **Fig. 1j, 1m**, 1040 nm, 65% excitation in **Fig. S2b**) and suboptimal wavelengths (940 nm, 10 % excitation in **Fig. S2b**) due to the high concentrations of circulating Alb-mSc-ST. Images were acquired using ThorImage Software Version 3.0. The laser power under the objective lens was measured by a power meter (Thorlabs) before imaging. The excitation power was linearly scaled with imaging depth (5−30 mW for 0−200 μm, 5−35 mW for 0−500 μm, 5−40 mW for 0−1000 μm) for volumetric imaging (field of view: ∼ 500 x 500 µm for **Fig. 1j−l**, **2e**, 250 x 250 µm for **Fig. 1m**), whereas it was kept around 14-20 mW for small volumetric imaging (field of view: ∼ 240 x 240 µm for **Fig. 2g, S2f**, ∼ 120 x 120 µm for **Fig. 3d**) and for time-lapse imaging of capillaries, arterioles and venules (field of view: ∼ 240 x 240 µm for **Fig. 2j—m, S2g−o**). Only for SC-GINKO experiments, the power was kept around 30 mW for time-lapse imaging (field of view: ∼ 500 x 500 µm for **Fig. 2h−i, S2q−r**) due to the low baseline GINKO2 signals.

For testing SC-iLACCO (**Fig. 2g, S2f**), awake mice were intraperitoneally injected with lactate (Sigma, 71718, 2 mg/g) and small volumetric images were subsequently acquired before injection and at 10 min, 20 min, 30 min, 40 min post-injection.

For whisker stimulation in anesthetized (70 mg/kg ketamine, 10 mg/kg xylazine) knock-in mice expressing SC-SEP or SC-iLACCO or SC-mNG (**Fig. 2j−m, S2g−o**), the left whiskers were stimulated by air puff (100-ms-long pulses, 5 Hz, 20 psi, duration of 2 min). The response of arterioles and venules was recorded by time-lapse imaging. Each recording session consisted of 30 s baseline, whisker stimulation and ∼6 min post-stimulation. Additionally, SC-iLACCO was also recorded in absence of stimulation (**Fig. S2m−o**).

All SC-GINKO experiments were performed in anesthetized (70 mg/kg ketamine, 10 mg/kg xylazine) mice. For testing SC-GINKO, either saline or artificial CSF containing 1 M KCl were locally applied to the second acute open craniotomy (**Fig. 2h, S2q**). Three time-lapse imaging were performed on the primary craniotomy: a 2-min baseline recording with saline, a 10-min recording with 1 M KCl and, after 3 washes with saline, a 2-min recording with saline. For the cortical spreading depression induction (**Fig. 2i, S2r**), neuronal ChR2 was optogenetically stimulated^25^ by a 473 nm blue laser (MBL-FN-473-200mW, 10 mW for 10s) in the second chronic craniotomy in the contralateral hemisphere. Two time-lapse imaging were performed on the primary craniotomy: a 2-min baseline recording and, after optogenetic stimulation, a 45-min recording.

For experiments inducing Alb-mSc-ST extravasation *in vivo* (**Fig. 3d**), a vessel was first identified in anesthetized knock-in mice expressing astrocytic ECS-SC or ECS-dSC and the BBB was then temporarily impaired by focused high-power laser irradiation (90**−**150mW for few seconds)^17^. Small volumetric images were taken before and after laser irradiation.

### *Ex Vivo* One-Photon Confocal Imaging

For intravital imaging of the colon or small intestine, mice were anaesthetized, their abdomens were incised, and their colon or small intestine was aspirated and fixed with a mouse adsorption fixer (Olympus, Tokyo, Japan) under 1.5% isoflurane. Alb-mSc-ST fluorescence in the colon or small intestine was then observed using an upright microscope (BX51WI, Olympus, Tokyo, Japan) with a 40x water immersion objective (LUMPLFLN40XW; NA = 0. 8; Olympus), a digital CMOS camera (C14440-20UP, Orca-Fusion, Hamamatsu Photonics, Shizuoka, Japan), and a Nipkow disc-type confocal scanning unit (CSU-W1, Yokogawa electric corporation, Tokyo, Japan) equipped with an optical pumped semiconductor laser (561 nm; 50 mW; Sapphire 561LP, Coherent). A custom-made filter set (CSU-DM/EM-SETSP66, Ex 562/30, Em 617/73, Yokogawa electric corporation) was used to detect the red fluorescence of mScarlet. The microscope was controlled and photographed using MetaMorph (version 7.8.13, Molecular devices, CA, USA), and time-lapse images were taken every 500 ms for 60 s.

### GRIN Lens Implantation and Baseplate Fixation for One-Photon Miniscope Imaging

Mice were anesthetized with 4–5% isoflurane for induction, 1–2% for maintenance (Vetflurane, Virbac), and secured in a stereotaxic frame (Kopf). Body temperature was maintained at 37°C using a heating pad. Ocrigel ointment was applied to the eyes to prevent dryness. The non-steroidal anti-inflammatory agent meloxicam was injected intraperitoneally to prevent post-surgical pain (Metacam, 4 mg/kg). The local anesthetic lidocaine (5 mg/kg) was given subcutaneously, prior to performing an incision of the skin in the anterior-posterior axis to expose the skull.

For vascular and neuronal imaging in prefrontal cortex (PFC) (**Suppl. Video 6**), a craniotomy was drilled above the right prefrontal cortex (2.0 mm anterio-posterior; +0.6 mm medio-lateral relative to bregma). AAV5/CamKII-GCaMP6f-WPRE-SV40 (2.3 x 10^13^ vg/mL; Addgene #100834-AAV) was unilaterally injected into the PFC (300 nL) by pressure application using pulled glass pipettes (tip diameter ∼ 20 μm) at the same coordinates and at a depth of 1.4 mm relative to brain surface. Five minutes after injection, the pipette was slowly retracted and a graded index lens (GRIN lens, 0.6 × 7.3 mm, Inscopix) was inserted at a depth of 1.5 mm.

For NAC vessel imaging (**Suppl. Video 5**), a craniotomy was drilled above the right nucleus accumbens (+1.0 mm anterio-posterior; +1.9 mm medio-lateral relative to bregma) to allow for the slow insertion of a dual-color graded index lens (GRIN lens, 7 × 0.6 mm, Inscopix) at the same coordinates and at a depth of 4.9 mm relative to brain surface.

The GRIN lens was then secured in place using dental cement (Super-Bond, Sun medical) and covered with silicon (Kwik-Sil, World Precision Instruments). After the surgery, mice were allowed to recover for 2 weeks, and postoperative monitoring was ensured for a minimum of 3 days.

Three weeks after surgery, mice were anaesthetized with isoflurane (4–5% for induction, 1–2% for maintenance) to fix the miniature microscope baseplate on top of the cranium implant (blue-light curable glue; Vertise Flow, Kerr), under visual control of the focal plane. The miniature microscope was then detached, the baseplate was capped with a baseplate cover (Inscopix) and the mouse was returned to its home cage.

### One-Photon Miniscope Imaging

Prior to recordings, mice were habituated to restraint and miniscope mounting. A dual-colour miniature microscope (nVue, Inscopix) was mounted immediately before each imaging session. Images were collected using nVue HD software (Inscopix Inc.). Data were acquired at a frame rate of 100 Hz. For PFC dual-colour recordings, green (GCaMP6f) and red (Alb-mScarlet) were acquired in an interleaved manner with a 1:4 ratio. Miniscope settings were as followed: for red channel, 545–560 nm LED, power of 1-2 mW/mm^2^ (excitation irradiance at objective front surface), acquisition bandpass 600–660 nm, analog gain of 4 to 8; for green channel, 455-480 nm LED, power of 1.5-2 mW/mm^2^, acquisition bandpass 499–530 nm; field of view of 640 x 400 pixels window (about 512 x 320 µm, nVue, Inscopix).

### Mannitol Injection

For experiments with mannitol (**Fig. 3f**), a 20% mannitol solution was made by dissolving 20 g of Mannitol powder (D-Mannitol Sigma-Aldrich M4125-100G) in 100 mL preheated (50°C) distilled water. After adjusting pH to ∼ 7, the solution was filtered through 0.22 µm syringe filter. Mice were injected via retro-orbital sinus with 20% mannitol (1 mg/g) 40 min before perfusion.

### Image Data Analysis

#### All images were pre-processed using Image J

For *in vitro* comparison of ECS-SC and ECS-dSC (**Fig. 3b**), the highest 5th percentile of pixel intensities was considered as membrane-expressed signal, accounting for variations in cell confluency and transfection efficiency across different culture wells. The mean of these signals was computed to represent the signal for each image.

For *ex vivo* macro fluorescence imaging of blood samples (**Fig. 1d−f, S1d, 2c, 2f, S2a, S2c−d**), fluorescence was quantified by drawing a ROI covering the entire blood sample and subtracting the endogenous blood fluorescence measured in an un-injected (naïve) mouse. For mice expressing both Alb-mSc-ST and green biosensors, the ratio between green and red fluorescence was evaluated to normalize the biosensor signal over the amount of Alb-mScarlet present in the sample.

For two-photon images, motion-correction was applied using the ImageJ tool TurboReg^26^ when needed. For whisker stimulation experiments with SC-SEP or SC-iLACCO or SC-mNG (**Fig. 2j−m, S2g−o**), ROIs within arterioles and venules of interest were drawn and the ratio between green fluorescence (SEP) and red fluorescence (mScarlet) was quantified so that to normalize the biosensor signal over the amount of flowing blood. The fluorescence ratio was then normalized to baseline (F−F_0_). Sensory-evoked functional hyperemia was assessed by measuring the vessel diameter. ROI was drawn perpendicularly to the vessel and the mean pixel intensity was plotted over time to create a kymograph. Vessel borders were defined using the full width half max function and their distance was measured to compute the arteriole diameter. The relative diameter change was normalized to baseline (F−F_0_) and smoothened by a mean filter (n = 20). For SC-iLACCO experiments with lactate injection (**Fig. 2g, S2f**), the average intensity of each volumetric image was projected and ROIs within vessels of interest were drawn. Similarly as before, the ratio between green fluorescence and red fluorescence was quantified and normalized to baseline (F−F_0_).

For *in vivo* validation of ECS-dSC using histological sections of brain that underwent capillary rupture by focused laser irradiation (**Fig. 3d**), 20 astrocytic domains were selected from the peri-hemorrhaged area and compared against similarly chosen domains in the contralateral homotopic site in 3 mice.

For histological validation of ECS-dSC using brain sections of mice injected with either mannitol or saline (**Fig. 3f**), two ROIs were drawn in the size of 0.3 mm x 0.3 mm from cortex and striatum for each animal. The entropy of selected ROIs was calculated using the Matlab function entropy^27^.

For the miniscope recordings (**Suppl. Videos 4 and 5**), data processing was conducted with IDPS software (Inscopix), using motion correction and (ΔF−F_0_) transformation for GCaMP signal. No spatial or temporal down sampling was applied. Furthermore, we applied a Gaussian Blur 3D filter (standard deviation: 2 pixels for x and y, 1 pixel for z) to **Suppl. Video 4** and we used *SUPPORT* to remove Poisson–Gaussian noise^28^ from the green channel in **Suppl. Video 5**.

### Statistical Analysis

All values are indicated as mean ± SEM. Comparison of two sample group means was assessed by t-test (parametric) or Mann Whitney test (non-parametric), as appropriate. Multiple group comparisons were performed using the following tests, as appropriate: two-way ANOVA or mixed-effects analysis using a REML model. Graph Prism 10 was used for all statistical analyses.

## Supporting information

Suppelmentary discussion

## Acknowledgements

This work was supported by KAKENHI grants (21H05614, YH; 22K06454/24H01221, AK; 22KK0140, MT; 24K01621, YN), the Novo Nordisk Foundation (NNFOC0058058, HaH), Danmarks Frie Forskningsfond (0134-00107B, HaH; 4285-00151B, AA), the Lundbeck Foundation (R360-2021-613, R436-2023-1219, HaH), National Institute of Health (U19NS128613, HaH, MaiN), the Naito Foundation (MasN), Nanken Kyoten (2021-kokusai 11, YH), AMED Brain/MINDS (JP21dm0207111, HiH), AMED Brain/MINDS 2.0 (JP24wm0625103, HiH), Marie Skłodowska-Curie Fellowship ANCoDy (101064009, AA), the Japan Science and Technology Agency (JPMJPR22E9, JPMJCR24T6, YN), Academia Sinica intramural funds (YN) and ONO Rising Star Fellowship (AA). We thank members of the laboratory for comments on earlier versions of the manuscript.

---

**Figure S1.**
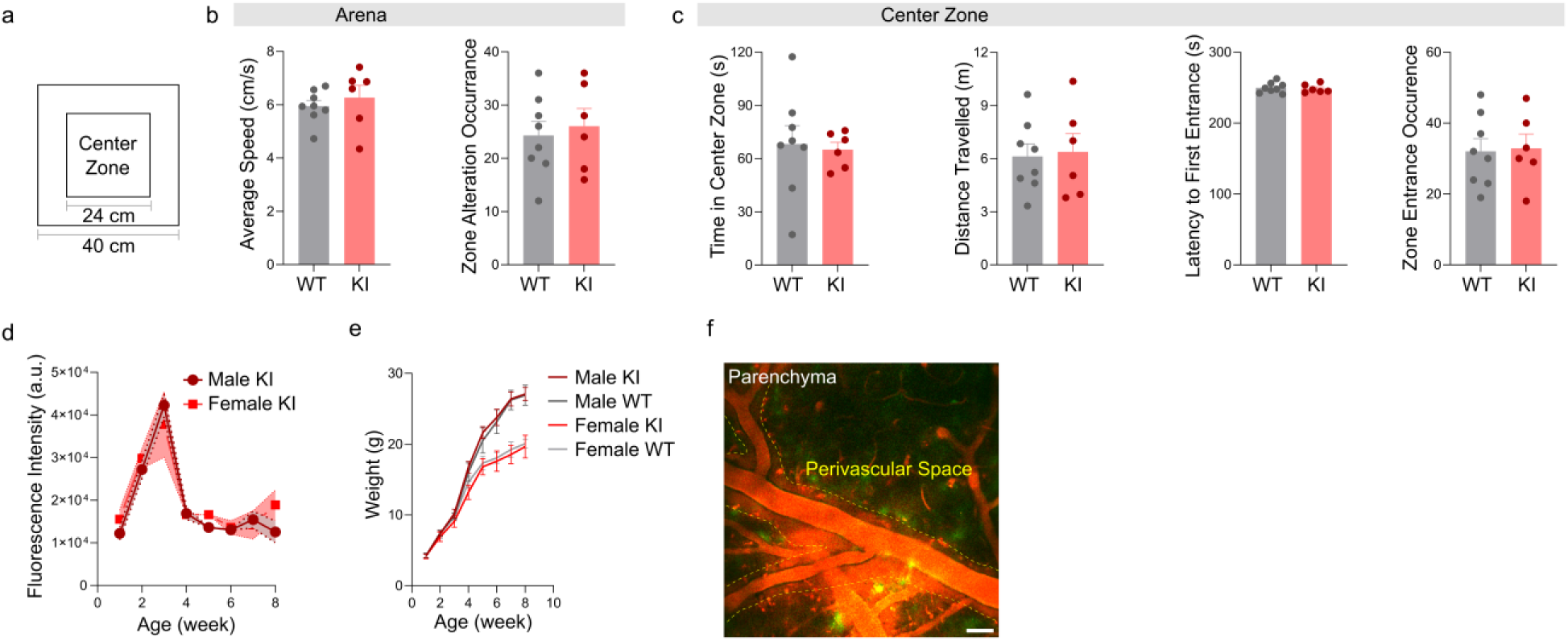
Extended data from Figure 1. (a) Schematic of the arena used for open field test. (b) Average speed (left) and zone alteration occurrence (right) in the arena for the last 6 min of 10 min recording (n = 6−8 mice). Unpaired t-test: p > 0.05. (c) From left to right: time in center zone, distance travelled, latency to first entrance and zone entrance occurrence in the center zone for the last 6 min of 10 min recording (n = 6−8 mice). Unpaired t-test: p > 0.05. (d) Fluorescence intensity in blood samples from male and females KI littermates in the first 8 weeks after birth (n = 3 mice). Two-way ANOVA: no significant effect of sex (F_1, 4_ = 0.6132, p = 0.4774). (e) Weight measured in KI and WT littermates in the first 8 weeks after birth (n = 3−4 mice per group). Two-way ANOVA: no significant effect of genotype (F_1, 5_ = 0.2591, p = 0.6324 for males; F_1, 5_ = 1.046, p = 0.3534 for females). (f) Representative image of pial vessels with labeled perivascular space from an Alb-mSc-ST x Prox1-EGFP^+^ mouse. Scale bar, 50 µm. All graphs show means ± SEM.

**Figure S2.**
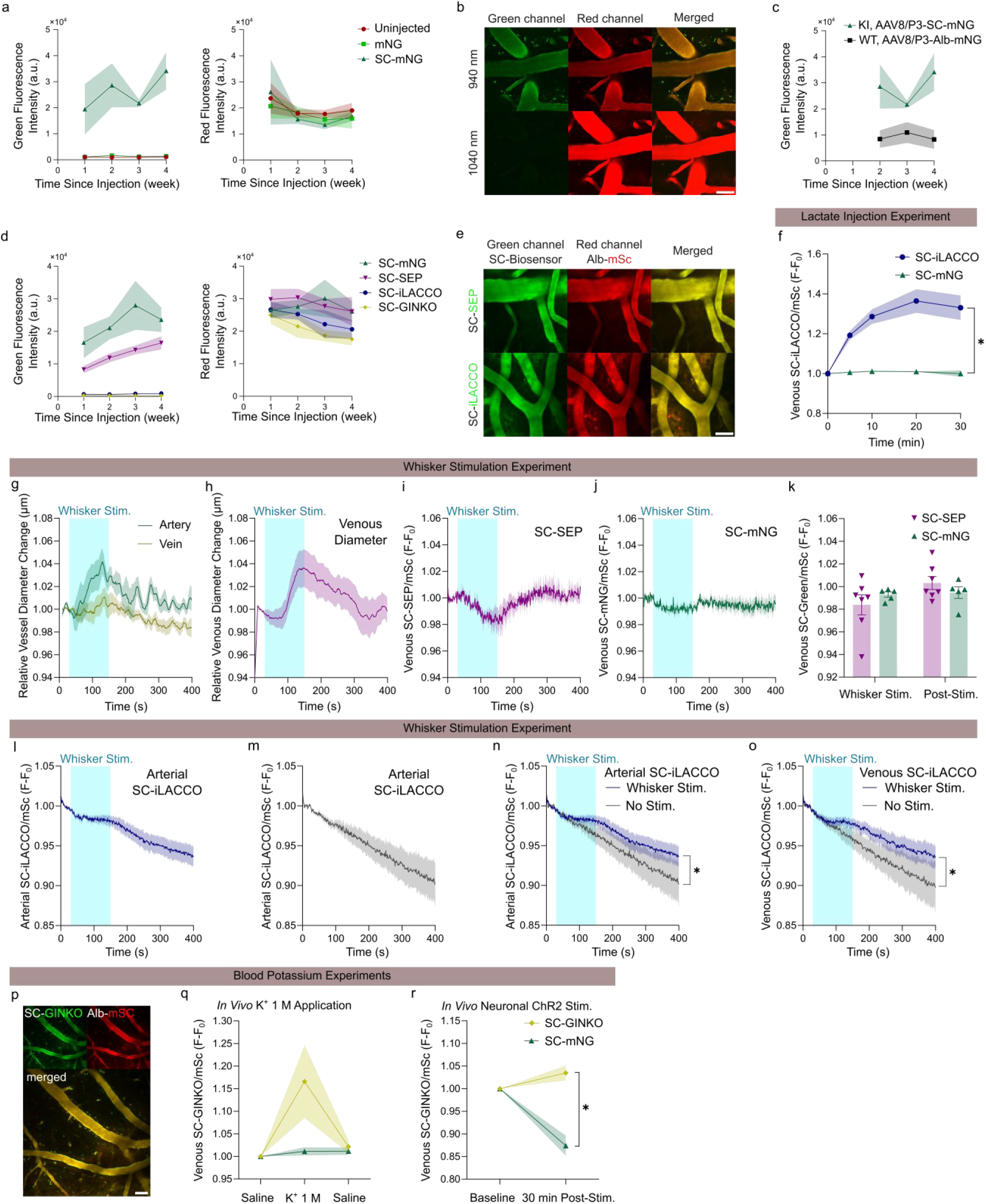
Extended data from Figure 2. (a) Green (left) and red fluorescence intensities (right) in blood samples collected from Alb-mSc-ST mice injected with AAV8/P3-mNG or AAV8/P3-SC-mNG or uninjected weekly in the first 4 weeks post-injection (n = 4−5 mice). (b) Two-photon images showing cortical vasculature of Alb-mSc-ST mice imaged at 940 nm (up) or 1040nm (bottom). Scale bar, 50 µm. (c) Green fluorescence intensity in blood samples collected from Alb-mSc-ST mice injected with AAV8/P3-SC-mNG and WT littermates injected with AAV8/P3-Alb-mNG (n = 3−5 mice). Two-way ANOVA: no significant effect of AAV type (F_1, 6_ = 5.814, p = 0.0525). (d) Green (left) and red fluorescence intensities (right) in blood samples collected from Alb-mSc-ST mice injected with AAV8/P3-SC-mNG or AAV8/P3-SC-SEP or AAV8/P3-SC-iLACCO weekly in the first 4 weeks post-injection (n = 5−7 mice). (e) Two-photon images showing cortical vasculature of Alb-mSc-ST mice injected with AAV8/P3-SC-SEP (top) or AAV8/P3-SC-iLACCO (bottom). Scale bar, 50 µm. (f) Green/red signal ratios in cortical venules of awake Alb-mSc-ST mice injected with AAV8/P3-SC-iLACCO or AAV8/P3-SC-mNG and imaged after intraperitoneal injection of lactate (2 mg/g, n = 5−6 mice). Two-way ANOVA: significant effect of AAV type (F_1, 9_ = 54.41, ****p < 0.0001). (g−k) Whisker stimulation experiment in anesthetized Alb-mSc-ST mice injected with AAV8/P3-SC-mNG, AAV8/P3-SC-SEP (n = 5−7 mice). Relative diameter change of vessels in mNG mice (g) and of venules in SEP mice (h). Green/red signal ratios in cortical venules of SEP and mNG mice (i−j). Quantification of normalized venous signals (green/red) during and after whisker stimulation (k). Two-way ANOVA: no significant effect of stimulation x AAV type interaction (F_1, 10_ = 1.281, p = 0.2841). (l−o) Whisker stimulation experiment in anesthetized Alb-mSc-ST mice injected with AAV8/P3-SC-iLACCO (n = 4−7 mice). Green/red signal ratio in cortical arterioles during whisker stimulation (l) or in absence of stimulation (m). Signal comparison with or without whisker stimulation in cortical arterioles (n) or venules (o). Two-way ANOVA: significant effect of time (F_1.376, 12.38_ = 29.82, ****p < 0.0001 for arterioles, F_1.290, 11.61_ = 26.55, ***p = 0.0001 for venules) and of stimulation x time interaction (F_399, 3591_ = 1.455, ****p < 0.0001 for arterioles, F_399, 3591_ = 1.707, ****p < 0.0001 for venules). (p) Two-photon images showing cortical vasculature of Alb-mSc-ST mice injected with AAV8/P3-SC-GINKO. Scale bar, 50 µm. (q) Green/red signal ratios in cortical venules of anesthetized Alb-mSc-ST mice injected with AAV8/P3-SC-GINKO or AAV8/P3-SC-mNG following application of either saline or 1 M KCl to a distant cortical site (n = 1−2 recordings from 2−3 mice). Two-way ANOVA: significant effect of applied solution x AAV type interaction (F_2,8_ = 3.543, p = 0.0791). (r) Green/red signal ratios in cortical venules of anesthetized Alb-mSc-ST mice injected with AAV8/P3-SC-GINKO or AAV8/P3-SC-mNG following optogenetically induced CSD (n = 1−2 recordings from 2−3 mice). Two-way ANOVA: significant effect of time x AAV type interaction (F_1, 3_ = 34.89, p = 0.0097). All graphs show means ± SEM.

**Figure S3.**
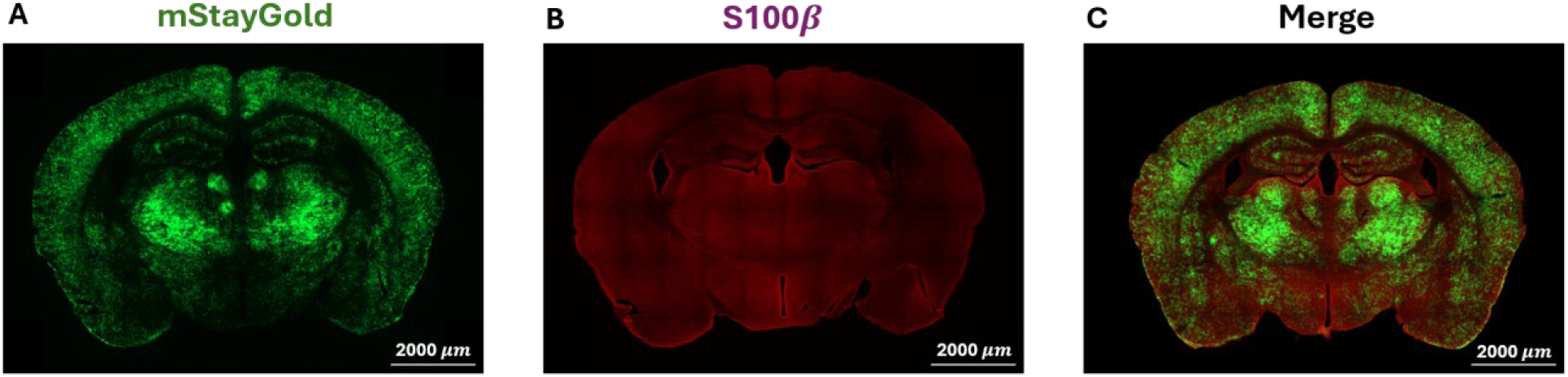
Representative overview of mStayGold and S100β signal distribution across a coronal brain section. (A) mStayGold reporter signal shown in green. Two AAV vectors were administered: AAV-PHP.eB/2-hGFAP-Cre-4×6(miRT)-hGHp(A) and AAV-PHP.eB/2-CAG-DIO-ECS-dSpyCatcher-mStayGold (each ∼3×10^11^ vg). (B) S100β immunolabelling shown in red. (C) Merged image of mStayGold and S100β channels. mStayGold signal was detectable across the section but appeared regionally variable rather than evenly distributed. We estimate that ∼70–80% of cortical areas are represented by ECS-dSC+ positive astrocytes. Scale bars: 2000 μm

## Supplementary Video 1

Supplementary Video 1. Flow in cortical capillary of an Alb-mSc-ST-mouse. Red blood cells appear as dark shadows against the white background provided by fluorescent albumin. Scale bar, 10 µm.

## Supplementary Video 2

Supplementary Video 2. Peripheral tissues of Alb-mSc-ST mice. One-photon macroscopic imaging of ear skin (left) and one-photon confocal imaging of colon and small intestine (center and right, respectively). Scale bar, 100 µm, 20 µm and 20 µm, respectively.

## Supplementary Video 3

Supplementary Video 3. Volumetric imaging (140 µm) of dural blood vessels (red) and lymph vessels (green) in an Alb-mSc-ST x Prox1-EGFP^+^ mouse. Scale bar, 10 µm.

## Supplementary Video 4

Supplementary Video 4. GRIN lens imaging of nucleus accumbens in an Alb-mSc-ST mouse. Scale bar, 50 µm.

## Supplementary Video 5

Supplementary Video 5. GRIN lens imaging of prefrontal cortex in an Alb-mSc-ST mouse injected with AAV5/CamKII-GCaMP6f. Scale bar, 50 µm.

## References

1. Iliff, J. J. et al. A paravascular pathway facilitates CSF flow through the brain parenchyma and the clearance of interstitial solutes, including amyloid β. Sci Transl Med 4, 147ra111 (2012).

2. Nedergaard, M. & Goldman, S. A. Glymphatic failure as a final common pathway to dementia. Science 370, 50–56 (2020).

3. Quinlan, G. J., Martin, G. S. & Evans, T. W. Albumin: Biochemical properties and therapeutic potential†. Hepatology 41, 1211–1219 (2005).

4. Zaias, J., Mineau, M., Cray, C., Yoon, D. & Altman, N. H. Reference Values for Serum Proteins of Common Laboratory Rodent Strains. J Am Assoc Lab Anim Sci 48, 387–390 (2009).

5. Habgood, M. D., Sedgwick, J. E., Dziegielewska, K. M. & Saunders, N. R. A developmentally regulated blood-cerebrospinal fluid transfer mechanism for albumin in immature rats. J Physiol 456, 181– 192 (1992).

6. Wang, X. et al. Liver-secreted fluorescent blood plasma markers enable chronic imaging of the microcirculation. Cell Reports Methods 2, 100302 (2022).

7. Vittani, M. et al. Virally Induced CRISPR/Cas9-Based Knock-In of Fluorescent Albumin Allows Long-Term Visualization of Cerebral Circulation in Infant and Adult Mice. in Fluorescence Imaging of the Brain (ed. Rusakov, D.) 127–144 (Springer US, New York, NY, 2024). doi:10.1007/978-1-0716-4011-1_6.

8. Bindels, D. S. et al. mScarlet: a bright monomeric red fluorescent protein for cellular imaging. Nat Methods 14, 53–56 (2017).

9. Keeble, A. H. et al. Approaching infinite affinity through engineering of peptide–protein interaction. Proceedings of the National Academy of Sciences 116, 26523–26533 (2019).

10. Møllgård, K. et al. A mesothelium divides the subarachnoid space into functional compartments. Science 379, 84–88 (2023).

11. Sankaranarayanan, S., De Angelis, D., Rothman, J. E. & Ryan, T. A. The use of pHluorins for optical measurements of presynaptic activity. Biophys J 79, 2199–2208 (2000).

12. Steiert, F., Petrov, E. P., Schultz, P., Schwille, P. & Weidemann, T. Photophysical Behavior of mNeonGreen, an Evolutionarily Distant Green Fluorescent Protein. Biophysical Journal 114, 2419–2431 (2018).

13. Hario, S. et al. High-Performance Genetically Encoded Green Fluorescent Biosensors for Intracellular l-Lactate. ACS Cent. Sci. 10, 402–416 (2024).

14. Nasu, Y. et al. Lactate biosensors for spectrally and spatially multiplexed fluorescence imaging. Nat Commun 14, 6598 (2023).

15. Wu, S.-Y. et al. A sensitive and specific genetically-encoded potassium ion biosensor for in vivo applications across the tree of life. PLOS Biology 20, e3001772 (2022).

16. Li, X. et al. Generation of destabilized green fluorescent protein as a transcription reporter. J Biol Chem 273, 34970–34975 (1998).

17. Reeson, P., Boghozian, R., Cota, A. P. & Brown, C. E. Optical opening of the blood-brain barrier for targeted and ultra-sparse viral infection of cells in mouse cortex. Cell Reports Methods 3, 100489 (2023).

18. McCarty, D. M., DiRosario, J., Gulaid, K., Muenzer, J. & Fu, H. Mannitol-facilitated CNS entry of rAAV2 vector significantly delayed the neurological disease progression in MPS IIIB mice. Gene Ther 16, 1340–1352 (2009).

19. Fu, B. M. Tumor Metastasis in the Microcirculation. Adv Exp Med Biol 1097, 201–218 (2018).

20. Jin, K. A Microcirculatory Theory of Aging. Aging Dis 10, 676–683 (2019).

21. Wang, X. et al. Liver-secreted fluorescent blood plasma markers enable chronic imaging of the microcirculation. Cell Reports Methods 2, 100302 (2022).

22. Konno, A. & Hirai, H. Efficient whole brain transduction by systemic infusion of minimally purified AAV-PHP.eB. Journal of Neuroscience Methods 346, 108914 (2020).

23. Yardeni, T., Eckhaus, M., Morris, H. D., Huizing, M. & Hoogstraten-Miller, S. Retro-orbital injections in mice. Lab Anim 40, 155–160 (2011).

24. Terunuma, M. et al. Postsynaptic GABAB receptor activity regulates excitatory neuronal architecture and spatial memory. J Neurosci 34, 804–816 (2014).

25. Houben, T. et al. Optogenetic induction of cortical spreading depression in anesthetized and freely behaving mice. J Cereb Blood Flow Metab 37, 1641–1655 (2017).

26. Thevenaz, P., Ruttimann, U. E. & Unser, M. A pyramid approach to subpixel registration based on intensity. IEEE Transactions on Image Processing 7, 27–41 (1998).

27. Gonzalez, R. C., Woods, R. E. & Eddins, S. L. Digital Image Processing Using MATLAB. (Prentice-Hall, Inc., USA, 2003).

28. Eom, M. et al. Statistically unbiased prediction enables accurate denoising of voltage imaging data. Nat Methods 20, 1581–1592 (2023).

