## Supplementary material for "Functional and structural profiling of circulation via genetically encoded modular fluorescent probes": Suppelmentary discussion

### Supplementary Discussion

We introduced a novel transgenic mouse model expressing mScarlet-fused albumin, thereby permanently labeling blood and other body fluids. Beyond an increase in brightness by fifty fold, (**Fig. 1d)**, this mouse overcomes several limitations of our previously developed AAV-based methods[1,2], such as the sex-dependent tropism of AAV8 to hepatocytes[3], feasibility of early-stage neonate imaging, and inefficient AAV re-administrations due to immune reactions[4]. Moreover, the mScarlet red fluorescent signal faithfully reflects the plasma albumin level since the expression originates from the genomic albumin locus. We observed plasma fluorescence peaking at around 3 weeks of age (**Fig. 1f)**, similar to reported albumin mRNA expression which increases from birth and peaks around 4 weeks in mice[5]. The vascular organization of the liver matures at around 3−4 weeks of age[5], thus allowing for more albumin to be released into the blood stream. After 4 weeks, plasma albumin decreases and stabilizes at a plateau concentration (**Fig. 1f**). This decrease is presumably due to blood volume increasing faster than that of liver in the first postnatal weeks, with neonates having slightly higher blood volume per unit of body weight than adult mice[6,7]. The Alb-mSc-ST enables to track such longitudinal concentration changes by fluorescence imaging. Of note, this transgenic mouse was successfully used with fiber photometry by our group to show norepinephrine-mediated vasomotion as a driver of cerebral spinal fluid dynamics[8].

Alb-mSc-ST mice offer an optimal solution for in-depth characterization of brain vasculature. Skull and dural blood vessels appear leaky to albumin (**Fig. 1k**), as manifested by high mScarlet signals in the interstitial fluid, similar to peripheral tissues (**Suppl. Video 2**). The origin and turnover of interstitial albumin such as in the dura mater is a subject of future studies. Interestingly, skull osteocytes and dural macrophages present red fluorescence, conceivably due to a combination of endogenous autofluorescence[9] and possible uptake of ECS Alb-mSc.

Notably, liquid compartments with lower albumin content, such as lymph (**Fig. 1l**) and interstitial fluid (**Fig. 1m**) were also visualized. As albumin extravasation from blood vessels is prevented by BBB in brain parenchyma, albumin in interstitial fluid is likely originated from CSF. Moreover, this mouse is readily available to study albumin-containing compartments in the periphery, as demonstrated by our recordings in skin, colon and small intestine (**Suppl. Video 2**). While we focused on heterozygous mice, homozygous mice are viable and could be used when even higher signals are needed.

Another utility of Alb-mSc-ST mice is investigation of the blood microenvironment via circulating biosensors through ST-SC bonding. We demonstrated the feasibility of plasma-targeted ST-SC-bonded macromolecules by *ex vivo* blood sample fluorescence and Western Blot (**Fig. 2c–d**). Green fluorescence from Alb-mSc-ST-SC-mNG-containing blood was three times brighter than a standard AAV-expressed Alb-mNG while both SC-mNG and Alb-mNG were expressed using the same dosage and the P3 promoter (**Fig. S2c**). The enhanced green signal is unlikely due to the smaller size of SC-mNG since a even smaller secretory plasma label IgKL-mNG was shown to have lower signals [1]. It is plausible that the large macromolecules are less susceptible to leakage or extrusion to other tissues. Our results indicate that the ST-SC macromolecule formation is advantageous for blood-targeted biosensors. Accordingly, we successfully implemented three different biosensors. Imaging the pH sensor SEP in vasculature of barrel cortex during whisker stimulation-induced functional hyperemia revealed a 2% decrease in fluorescence ratio (**Fig. 2k, S2i**), corresponding to a 0.03 decrease in blood pH, which is within the physiological pH range. The pH decrease was very limited thanks to the efficient carbonic acid-bicarbonate buffering system[10]. Thereafter, blood pH quickly returned to baseline. This transient blood acidification could be originated from a metabolically derived $CO_2$ and/or lactate production with subsequent vessel absorption. Indeed, a recent study showed that hypercapnia induces blood pH decrease[11]. Remarkably, lactate released by astrocytes following whisker stimulation[12] can result in decreased brain pH[13,14], which will then be buffered by blood, similarly to the blood acidification observed concomitantly to lactate increase during exercise[15]. Of note, previous pH electrode measurements showed extracellular alkalization after - but not during - electrical forepaw stimulation in forelimb region of somatosensory cortex of anesthetized mice[16], whereas we observed whisker stimulation-induced blood acidification. This discrepancy might be due to the more intense stimulation paradigm in our experiment and targeting barrel cortex, where energy metabolism is high as manifested by cytochrome c oxidase activity[17].

Signal for the lactate biosensor iLACCO increases as lactate is absorbed by peripheral vessels after being intraperitoneally injected (**Fig. 2g, S2f**). Recordings in ketamine-xylazine anesthetized mice revealed a constant blood lactate decrease (**Fig. S2m–o**), likely due to a combination of metabolic suppression, reduced glycolytic activity and altered circulatory dynamics during aneasthesia[18]. The reduced rate of lactate reduction during whisker stimulation with a small delay (**Fig. S2l–o**) could be attributed to astrocyte-derived lactate[12].

Finally, we tested the potassium sensor GINKO. Blood potassium levels could be reversibly affected by addition/removal of artificial CSF containing high potassium to a second acute craniotomy (**Fig. 2h, S2q**). Excess increase of local extracellular potassium induces cortical spreading depression (CSD)[19]. Unfortunately, the CSD-induced potassium released from neurons and astrocytes and its potential subsequent vessel incorporation was masked with this paradigm because of the applied potassium saturating blood vessels. However, in the optogenetically induced CSD[20], an increase in blood potassium levels was indeed recorded several minutes after stimulation (**Fig. 2i, S2r**), indicating vasculature clearance as a route for excretion of the excess potassium released during the CSD event.

The Alb-mSc-ST mouse model can also be used for monitoring BBB permeability when combined with astrocyte-targeted AAV expressing SC in the ECS. BBB leakage is usually assessed *ex vivo* by dye injection (e.g. Evans Blue) followed by evaluation of leaked signal in the perfused brain[21] or *in vivo* macroscopically by MRI and PET and microscopically by two-photon microscopy with fluorescent dyes[22]. The latter offers the possibility to investigate the permeability of individual vessels, but smaller and slower BBB disruption events might remain undetectable as the extravasated blood elements diffuse in the ECS. Our method solves this issue by allowing astrocytes to capture the extravasated Alb-mSc-ST via ECS-SC binding. Indeed, we used laser irradiation[23,24] and systemic injection of the transiently permeabilizing mannitol[25] to temporarily disrupt BBB and we successfully observed the extravasated Alb-mSc-ST via ECS-SC binding *in vivo*, respectively (**Fig. 3d–f** ). With the current design, ECS-SC is a membrane protein subjected to lateral diffusion: consequently, Alb-mSc-ST bound to ECS-SC will label the entire astrocyte, despite the binding happening only at the level of astrocytic endfeet enveloping cerebral vessels. This could prove advantageous as the diffusing fluorescent signal might allow imaging of signals which would remain undetected if localized in endfeet and below the spatial resolution limit of the imaging technique. Conversely, for imaging techniques with better spatial resolution, future design should incorporate local targeting of endfeet. This novel assay represents the perfect tool for *in vivo* evaluation of functional or pathological BBB permeability, such as in activity-dependent plasticity[26] and neuropsychiatric disease[27], respectively.

Overall, we have presented a novel and multipurpose transgenic mouse model that will provide a resourceful platform for the investigation of body fluids including blood, cerebral extracellular fluid compartments, BBB permeability, and the blood plasma microenvironment.

#### Supplementary References


1. Wang, X. *et al.* Liver-secreted fluorescent blood plasma markers enable chronic imaging of the microcirculation. *Cell Reports Methods* **2**, 100302 (2022).

2. Vittani, M. *et al.* Virally Induced CRISPR/Cas9-Based Knock-In of Fluorescent Albumin Allows Long-Term Visualization of Cerebral Circulation in Infant and Adult Mice. in *Fluorescence Imaging of the Brain* (ed. Rusakov, D.) 127–144 (Springer US, New York, NY, 2024). doi:10.1007/978-1-0716-4011-1_6.

3. Piechnik, M. *et al.* Sex Difference Leads to Differential Gene Expression Patterns and Therapeutic Efficacy in Mucopolysaccharidosis IVA Murine Model Receiving AAV8 Gene Therapy. *International Journal of Molecular Sciences* **23**, 12693 (2022).

4. Shinohara, Y. *et al.* Effects of Neutralizing Antibody Production on AAV-PHP.B-Mediated Transduction of the Mouse Central Nervous System. *Mol Neurobiol* **56**, 4203–4214 (2019).

5. Nakagaki, B. N. *et al.* Immune and metabolic shifts during neonatal development reprogram liver identity and function. *Journal of Hepatology* **69**, 1294–1307 (2018).

6. Moreno-Carranza, B. *et al.* Prolactin regulates liver growth during postnatal development in mice. *Am J Physiol Regul Integr Comp Physiol* **314**, R902–R908 (2018).

7. Liu, Z.-J. *et al.* Expansion of the neonatal platelet mass is achieved via an extension of platelet lifespan. *Blood* **123**, 3381–3389 (2014).

8. Hauglund, N. L. *et al.* Norepinephrine-mediated slow vasomotion drives glymphatic clearance during sleep. *Cell* **188**, 606-622.e17 (2025).

9. Di Guardo, G. Lipofuscin, Lipofuscin-Like Pigments and Autofluorescence. *Eur J Histochem* **59**, 2485 (2015).

10. Atkinson, D. E. & Bourke, E. Metabolic aspects of the regulation of systemic pH. *American Journal of Physiology-Renal Physiology* **252**, F947–F956 (1987).

11. Tournissac, M. *et al.* Neurovascular coupling and CO2 interrogate distinct vascular regulations. *Nat Commun* **15**, 7635 (2024).

12. Magistretti, P. J. & Allaman, I. Lactate in the brain: from metabolic end-product to signalling molecule. *Nat Rev Neurosci* **19**, 235–249 (2018).

13. Prabakaran, S. *et al.* Mitochondrial dysfunction in schizophrenia: evidence for compromised brain metabolism and oxidative stress. *Mol Psychiatry* **9**, 684–697 (2004).

14. Hagihara, H. *et al.* Large-scale animal model study uncovers altered brain pH and lactate levels as a transdiagnostic endophenotype of neuropsychiatric disorders involving cognitive impairment. *eLife* **12**, RP89376 (2024).

15. Goto, K. *et al.* Hormonal and metabolic responses to slow movement resistance exercise with different durations of concentric and eccentric actions. *Eur J Appl Physiol* **106**, 731–739 (2009).

16. Theparambil, S. M. *et al.* Astrocytes regulate brain extracellular pH via a neuronal activity-dependent bicarbonate shuttle. *Nat Commun* **11**, 5073 (2020).

17. Wong-Riley, M. T. & Welt, C. Histochemical changes in cytochrome oxidase of cortical barrels after vibrissal removal in neonatal and adult mice. *Proceedings of the National Academy of Sciences* **77**, 2333–2337 (1980).

18. Ahmadi-Noorbakhsh, S. *et al.* Anesthesia and analgesia for common research models of adult mice. *Laboratory Animal Research* **38**, 40 (2022).

19. Somjen, G. G. Mechanisms of Spreading Depression and Hypoxic Spreading Depression-Like Depolarization. *Physiological Reviews* **81**, 1065–1096 (2001).

20. Houben, T. *et al.* Optogenetic induction of cortical spreading depression in anesthetized and freely behaving mice. *J Cereb Blood Flow Metab* **37**, 1641–1655 (2017).

21. Saunders, N. R., Dziegielewska, K. M., Møllgård, K. & Habgood, M. D. Markers for blood-brain barrier integrity: how appropriate is Evans blue in the twenty-first century and what are the alternatives? *Front. Neurosci.* **9**, (2015).

22. Harris, W. J. *et al.* In vivo methods for imaging blood–brain barrier function and dysfunction. *Eur J Nucl Med Mol Imaging* **50**, 1051–1083 (2023).

23. Reeson, P., Boghozian, R., Cota, A. P. & Brown, C. E. Optical opening of the blood-brain barrier for targeted and ultra-sparse viral infection of cells in mouse cortex. *Cell Reports Methods* **3**, 100489 (2023).

24. Nishimura, N. *et al.* Targeted insult to subsurface cortical blood vessels using ultrashort laser pulses: three models of stroke. *Nat Methods* **3**, 99–108 (2006).

25. McCarty, D. M., DiRosario, J., Gulaid, K., Muenzer, J. & Fu, H. Mannitol-facilitated CNS entry of rAAV2 vector significantly delayed the neurological disease progression in MPS IIIB mice. *Gene Ther* **16**, 1340–1352 (2009).

26. Swissa, E. *et al.* Cortical plasticity is associated with blood–brain barrier modulation. *eLife* **12**, RP89611 (2024).

27. Kealy, J., Greene, C. & Campbell, M. Blood-brain barrier regulation in psychiatric disorders. *Neurosci Lett* **726**, 133664 (2020).
